# Genome-wide association study of circadian rhythmicity in 71 500 UK Biobank participants and polygenic association with mood instability

**DOI:** 10.1101/350983

**Authors:** Amy Ferguson, Laura M. Lyall, Joey Ward, Rona J. Strawbridge, Breda Cullen, Nicholas Graham, Claire L. Niedzwiedz, Keira J.A. Johnston, Daniel MacKay, Stephany M. Biello, Jill P. Pell, Jonathan Cavanagh, Andrew M. McIntosh, Aiden Doherty, Mark E.S. Bailey, Donald M. Lyall, Cathy A. Wyse, Daniel J. Smith

## Abstract

**Background:** Circadian rhythms are fundamental to health and are particularly important for mental wellbeing. Disrupted rhythms of rest and activity are recognised as risk factors for major depressive disorder and bipolar disorder.

**Methods:** We conducted a genome-wide association study (GWAS) of low relative amplitude (RA), an objective measure of circadian rhythmicity derived from the accelerometer data of 71 500 UK Biobank participants. Polygenic risk scores (PRS) for low RA were used to investigate potential associations with psychiatric phenotypes.

**Outcomes:** Two independent genetic loci were associated with low RA, within genomic regions for Neurofascin (*NFASC*) and Solute Carrier Family 25 Member 17 (*SLC25A17*). A secondary GWAS of RA as a continuous measure identified a locus within Meis Homeobox 1 (*MEIS1*). There were no significant genetic correlations between low RA and any of the psychiatric phenotypes assessed. However, PRS for low RA was significantly associated with mood instability across multiple PRS thresholds (at PRS threshold 0·05: OR=1·02, 95% CI=1·01-1·02, p=9·6×10^−5^), and with major depressive disorder (at PRS threshold 0·1: OR=1·03, 95% CI=1·01-1·05, p=0·025) and neuroticism (at PRS threshold 0·5: Beta=0·02, 95% CI=0·007-0·04, p=0·021).

**Interpretation:** Overall, our findings contribute new knowledge on the complex genetic architecture of circadian rhythmicity and suggest a putative biological link between disrupted circadian function and mood disorder phenotypes, particularly mood instability, but also major depressive disorder and neuroticism.

## Introduction

Circadian rhythms are variations in physiology and behaviour that recur approximately every 24 hours.^1^ They include rhythms of body temperature, hormone release, activity, concentration, mood, eating and sleeping. Circadian rhythmicity is co-ordinated centrally by the suprachiasmatic nucleus in the anterior hypothalamus^2^ and plays a fundamental role in homeostasis and the maintenance of both physical and mental wellbeing.^2,3^ Disruption to circadian rhythmicity is associated with a range of adverse health outcomes, including cardiovascular disease, obesity, diabetes and some cancers,^4–6^ as well as increased risk for major depressive disorder (MDD) and bipolar disorder (BD).^7–9^

Circadian rhythmicity is regulated by both exogenous environmental stimuli (“zeitgebers”) and by genetic factors.^10^ Several core circadian clock genes are involved in autoregulatory transcription/translational feedback loops that maintain cell-cycle function.^11^ However, the control of circadian rhythms is likely to be polygenic, with regulatory genes and pathways still to be identified.^3,12^

To date, the most commonly used measure of circadian phenotypes has been subjectively-reported chronotype, defined as an individual’s preference for morning or evening wakefulness and activity.^13^ Evening chronotype is more likely to be associated with adverse health outcomes.^3,14–16^ Recently, genome wide association studies (GWAS) of chronotype, self-reported sleep duration, and accelerometer-derived sleep traits have identified several independent genetic loci previously implicated in the regulation of circadian function (including *PER2*, *PER3*, *RSG16*, *AK5*, *FBXL13*), in addition to novel associated genetic loci.^17–22^

However, as a subjective measure, chronotype can be vulnerable to response bias. It may also have inconsistent associations with more objective measures of circadian rhythmicity.^23^ In a recent study of over 91 488 individuals in the UK Biobank cohort, we derived objective measures of rest-activity rhythmicity from accelerometer data.^24^ We found that low relative amplitude (RA), a measure of an individual’s rest-activity rhythm, was associated with several mood disorder phenotypes.^24^ We now extend this work by conducting a GWAS of low RA in the largest sample known to date from UK Biobank. We also assess the degree of genetic correlation between low RA and several psychiatric phenotypes, including attention deficit hyperactivity disorder (ADHD), BD, MDD, mood instability, post-traumatic stress disorder (PTSD), schizophrenia, anxiety and insomnia. Further, within a group of UK Biobank participants who were not part of the primary GWAS study (up to 141 000 individuals), we test for association between increased polygenic risk score (PRS) for low RA and mood disorder phenotypes (specifically BD, MDD, generalised anxiety disorder (GAD), mood instability and neuroticism).

## Methods

### Participants and ethical approval

Over 502 000 United Kingdom (UK) residents aged 37-73 years were recruited to the UK Biobank cohort from 2006-2010. At one of 22 assessment centres across the UK, participants completed a range of lifestyle, demographic, health, mood, cognitive and physical assessments and questionnaires.^25^ Here, we used data from 91 448 participants who also provided accelerometer data that passed quality control (details below) and who had available data on all covariates included within fully and/or partially adjusted models for use in the GWAS. UK Biobank obtained informed consent from all participants and this study was conducted under generic approval from the NHS National Research Ethics Service (approval letter dated 13 May 2016, Ref 16/NW/0274) and under UK Biobank approvals for applications 12761 (PI Cathy Wyse; accelerometer data for use in GWAS) and 6553 (PI Daniel Smith; correlations with psychiatric traits).

### Accelerometry data collection and pre-processing

In 2013, 240 000 UK Biobank participants were invited to wear an accelerometer for seven days as part of a physical activity monitoring investigation.^26^ Of these, 103 720 (43%) accepted and returned the accelerometer to UK Biobank after use. Participants received a wrist-worn Axivity AX3 triaxial accelerometer in the post and were asked to wear the device on their dominant wrist continuously for seven days, while continuing with their normal activities. At the end of the seven-day period, participants were instructed to return the accelerometer to UK Biobank using a prepaid envelope. Accelerometers were calibrated to local gravity. Devices recorded data at a sampling rate of 94-104Hz, and data were resampled to 100Hz offline. Periods where no data were recorded for >1s were coded as missing, and machine noise was removed using a Butterworth low-pass filter (cut-off 20Hz). Raw activity intensity data were combined into five second epochs. Further details on data pre-processing^26^ are available from UK Biobank at http://biobank.ctsu.ox.ac.uk/crystal/refer.cgi?id=131600.

### Circadian rest-activity rhythmicity (relative amplitude)

From the summary five second epoch data, a measure of relative amplitude (RA) was calculated using Clocklab Version 6 (Actimetrics). This accelerometer-derived activity measure has demonstrated reliability and validity.^27^ RA is used commonly as a non-parametric measure of rest-activity rhythm amplitude. It is defined as the relative difference between the most active continuous 10-hour period (M10) and the least active continuous 5-hour period (L5) in an average 24-hour period, using the formula below:^28^

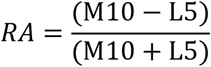

M10 is the mean activity during the continuous 10 hour period containing maximum activity in each 24 hour recording period (midnight to midnight). L5 is the mean activity for the corresponding 5 hour period containing the minimum activity within the same recording period. For each individual, the RA data point was the mean RA value across all included 24-hour periods (seven days). RA ranges from 0 to 1, with higher values indicating greater distinction between activity levels during the most and least active periods of the day.

Participants who provided accelerometer data for less than 72 hours or who did not provide data for all one-hour periods within the 24-hour cycle were excluded from analyses. Participants were also excluded if their data was identified by UK Biobank as having poor calibration. We also excluded over 10 000 participants with data flagged by UK Biobank as unreliable (unexpectedly small or large size) and participants whose wear-time overlapped with a daylight savings clock change.^24^ In the current sample (N=91,870), mean RA was 0·87 (SD=0·06; range 0·121-0·997), similar to previously reported values in healthy populations,^8^ however the distribution of RA values observed in our sample was negatively skewed (Supplemental Figure 1).

### Genotyping and imputation

UK Biobank released genotypic data for over 500 000 participants using two genotyping arrays specifically designed for UK Biobank with 95% shared marker content.^29^ Approximately 10% of these participants were genotyped using Applied Biosystems UK BiLEVE Axiom array by Affymetrix, with the remaining participants being genotyped using Applied Biosystems UK Biobank Axiom Array. Phasing on the autosomes was done using SHAPEIT3 using the 1000 Genomes Phase 3 dataset as a reference panel. Imputation of single nucleotide polymorphism (SNP) genotypes was carried out using IMPUTE4; the merged UK10K and 1000 Genomes Phase 3 reference panel, as used for the UK Biobank interim genotype data release. Stringent quality control was applied to the data, described in an open access document.^29^

### Primary GWAS of low relative amplitude

Our primary GWAS was a study of cases of low RA defined as cases with a mean RA greater than two standard deviations below the overall mean RA, with the remaining participants as controls.^24^ Before proceeding with genetic analyses, exclusions were applied to the data. Individuals were removed according to UK Biobank genomic analysis exclusions, failure of quality control, gender mismatch, sex chromosome aneuploidy, ethnicity (not Caucasian), lack of accelerometry data, plus other accelerometry exclusions, as noted above. For related individuals (first cousins or closer), a single individual was randomly selected from each pair of related individuals for inclusion in the analysis. After these (and the acclelerometry-based) exclusions, 71 500 individuals were available for GWAS. Data was further refined by removing SNPs with an imputation score of <0·8, minor allele frequency of <0·01 and Hardy-Weinberg equilibrium p<1×10^−6^ resulting in 7 969 123 variants remaining.

The primary association analysis was conducted using logistic regression in PLINK;^30^ an additive allelic effects model was used with sex, age, genotyping array, and the first eight genetic principal components as covariates. For the GWAS, genome-wide significance was less than p<5×10^−8^.

### Secondary GWAS of RA as a continuous trait

The BOLT-LMM method allows the inclusion of related individuals within GWAS. This method relaxes the assumptions of the standard GWAS, as used above, by using a mixture of two Gaussian priors and is a generalisation of a standard mixed model. This mixed model accounts for both relatedness and population stratification; this results in greater power when compared to principal component analysis.^31^ As above, individuals were removed according to UK Biobank genomic analysis exclusions, failure of quality control, gender mismatch, sex chromosome aneuploidy, ethnicity (not Caucasian), and lack of accelerometry data. After these exclusions, 77 440 individuals were available for this GWAS. As above, genome-wide significance was less than p<5×10^−8^. Note that due to the imbalance between cases and controls available for low RA, we were unable to use the BOLT-LMM approach for the primary GWAS.^31^

### Expression quantitative trait locus (eQTL) analysis

The lead SNP from each locus, identified by GWAS, was assessed for the possibility of expression quantitative trait loci (eQTLs). This genotype-specific gene expression was assessed using the GTEx portal. The portal was also used to investigate tissue-specific expression for the implicated genes.^32^

### Gene-based analysis

The summary statistics from both the primary and secondary GWAS analyses were uploaded to FUMA web application for gene-based analyses.^33^ Gene-based analyses were carried out based on the MAGMA method using all genetic associations within the summary statistics.^33,34^ For these analyses genome-wide significance was set at p<5×10^−5^.

### Genetic correlations between low RA and psychiatric phenotypes

Linkage Disequilibrium Score Regression (LDSR) was applied to the GWAS summary statistics to provide an estimate of SNP heritability (h^2^SNP).^35,36^ LDSR was also used to investigate genetic correlations between low RA and anxiety, ADHD, BD, MDD, mood instability, PTSD, schizophrenia and insomnia. The LD scores for these disorders were obtained using the summary statistics from the Psychiatric Genomics Consortium, CNCR-Complex Trait Genomics group, and UK Biobank. ADHD, BD, MDD, schizophrenia and insomnia were analysed using LD Hub.^36^

### Psychiatric diagnoses, neuroticism and mood instability

#### a) Bipolar Disorder, Major Depressive Disorder and Generalised Anxiety Disorder

A mental health questionnaire (MHQ) was developed by a mental health research reference group to collect additional mental health phenotype data in UK Biobank and was administered during 2016-2017.^37^ The MHQ was used to obtain information about individuals’ lifetime experiences of psychiatric disorders. The questions were based on a modified Composite International Diagnostic Interview Short Form (CIDI-SF). Lifetime depression (referred to here as ‘lifetime MDD’), ‘lifetime BD’ and lifetime generalised anxiety disorder (referred to as ‘lifetime GAD’) were evaluated based on answers provided by participants to the online MHQ. Therefore, these measurements represent a likelihood of diagnosis with the disorder of interest, rather than confirmed diagnoses.^37^ Individuals who self-reported BD or MDD were excluded from the control groups. The ‘lifetime’ variables were also mutually exclusive, designated cases for one variable were excluded from both the case and control groups for the other variables of interest. Due to these exclusions, the number of observations for each association tested was different.

#### b) Neuroticism

To define neuroticism a score was taken from the 12-item neuroticism scale of the Eysenck Personality Questionnaire-Revised Short Form (EPQ-R-S).^38,39^ Individuals were given a score of 0 or 1 for a “no or yes” answer to each item, with total score from 0 to 12.

#### c) Mood instability

A “mood instability” outcome measure was also obtained from the EPQ-R-S questionnaire: for one question participants were asked *“Does your mood often go up and down?”* and given the option to answer “yes”, “no”, “don’t know” or “prefer not to answer”^38^. Individuals who selected “don’t know” or “prefer not to answer” were coded as missing; this allowed the generation of a categorical mood instability variable where those who answered “yes” were designated as cases and participants who answered “no” were controls, those answering “don’t know” and “prefer not to answer” were excluded.^40^

### Association between PRS for low RA and affective disorder phenotypes

Associations between higher PRS for low RA and psychiatric diagnoses were examined in up to 76 018 individuals who had completed the MHQ and who were not included in the primary GWAS. Similarly, associations between low RA PRS and mood instability/neuroticism were examined in between 91 248 and 140 504 individuals (depending on the dependent variable) not included in the GWAS. PRS including SNPs at 6 different significance thresholds (p<5×10^−8^, p<5×10^−5^, p<0·01, p<0·05, p<0·1, p<0·5) were divided into quartiles, with the exception of p<5×10^−8^ which was divided into tertiles as only three different PRS were generated for this threshold. The top and bottom quantiles were compared in logistic regression models that were adjusted for age, sex, Townsend deprivation index,^41^ genotype array and the first eight genetic principal components. False discovery rate (FDR) correction was applied.^42,43^

## Results

### GWAS of low RA

Our primary analysis was a case-control GWAS of low RA. The GWAS data showed only a slight deviation in test statistics compared to the null (λ_GC_=1·016, Figure 1); this deviation may be due to the polygenic architecture of low relative amplitude. SNP heritability (h^2^_SNP_) accounted for less than 1% of the population variance (h^2^_SNP_=0·0067, se=0·0054). The Manhattan plot for low RA GWAS is presented in Figure 1. Two independent genomic loci on chromosomes 1 and 22 were associated with low RA at genome-wide significance, including three SNPs (described in Supplemental Table 1). These SNPs highlighted two candidate gene loci: Neurofascin (*NFASC*) on chromosome 1 and Solute Carrier Family 25 Member 17 (*SLC25A17*) on chromosome 22 (Supplemental Figure 2). As each of these SNPs is an intronic variant, the exact effect of each polymorphism is unclear.

**Figure 1.**
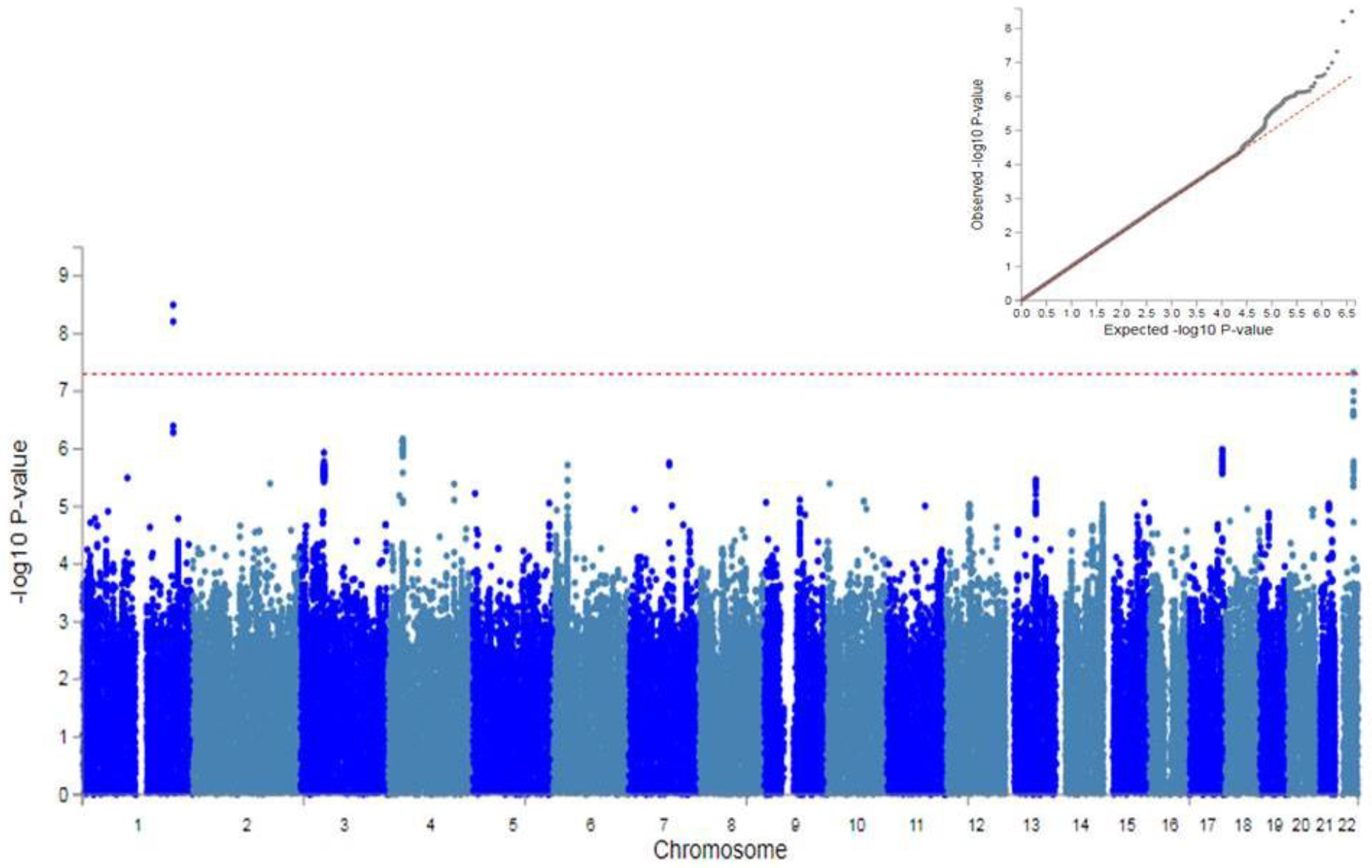
SNP Manhattan plot and QQ plot (inset) of low RA GWAS (N=2 700 cases verses N=68 300 controls) Red line of Manhattan plot represents genome-wide significance (p<5×10^−8^).

### GWAS of continuous RA

As a secondary analysis, we performed a GWAS of a continuous measure of RA using a BOLT-LMM model. The BOLT-LMM GWAS showed a slight deviation in the test statistics compared to the null (λ_GC_=1·054, Figure 2), again consistent with a polygenic architecture for RA. The h^2^_SNP_ for RA as a continuous measure accounted for greater than 8% of the population variance (h^2^_SNP_=0·085, se=0·00035). Five SNPs, all localised to one locus on chromosome 2, were associated with continuous RA at genome-wide significance (described in Supplemental Table 2). These SNPs highlight the Meis Homeobox 1 (*MEIS1*) gene. Again, as noted above, these are intronic SNPs and their exact effects are not known.

**Figure 2.**
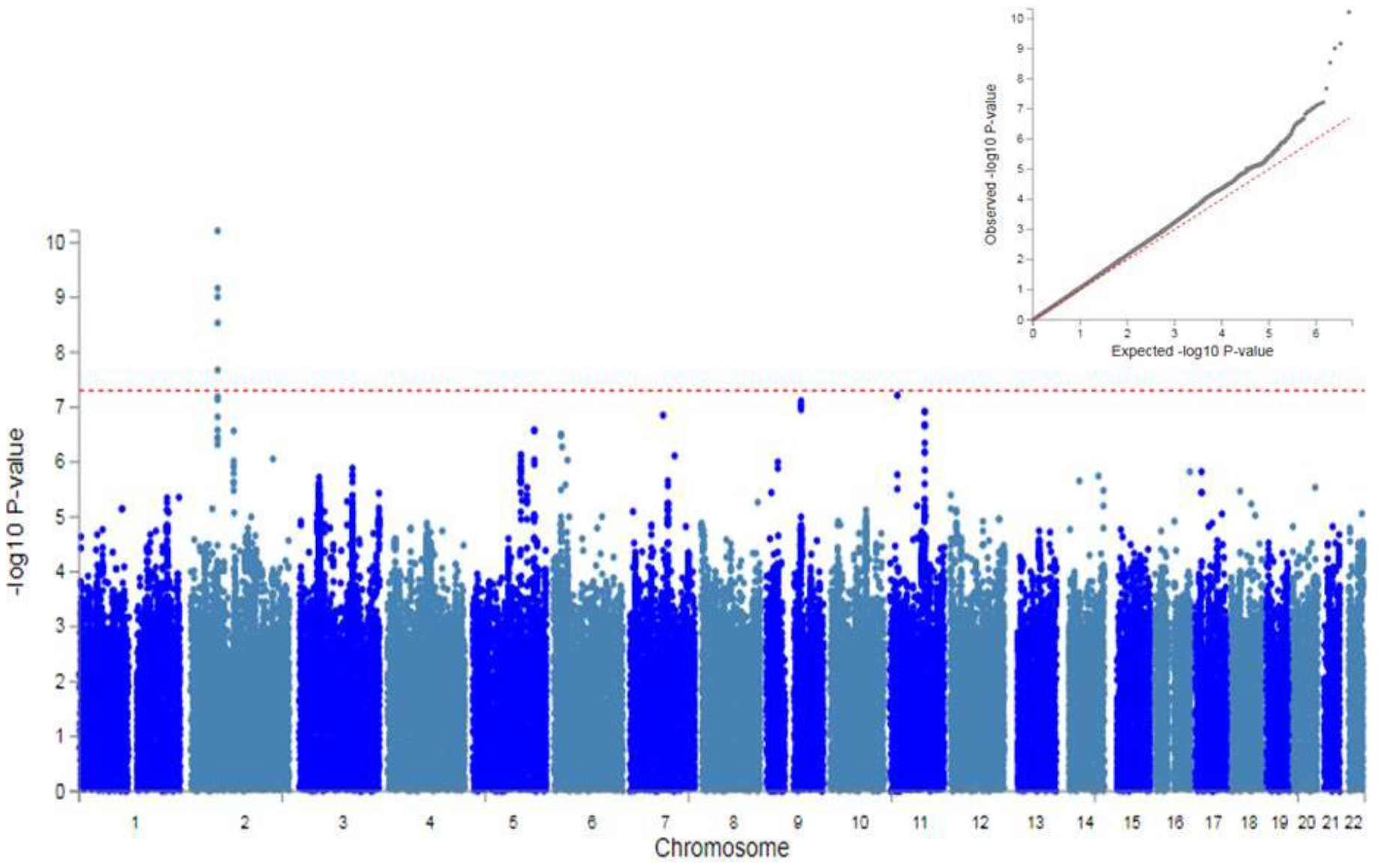
SNP Manhattan plot and QQ plot (inset) of continuous RA GWAS (N=77 440) Red line of Manhattan plot represents genome-wide significance (p<5×10^−8^).

### Expression quantitative trait loci (eQTL) analysis

The lead SNPs from both GWAS were assessed for potential eQTLs. Only the lead SNP found on chromosome 22 (rs9611417) was associated with the expression of a nearby gene. Being heterozygous at rs9611417 was associated with lower expression of *RANGAP1* gene in oesophageal mucosa in comparison to rs9611417 C allele homozygotes (Beta=-0·43, p=7·2×10^−5^, Supplemental Figure 5). Information on the influence of G homozygotes was not available.

### Gene-based analysis of RA

Gene-based analyses of both low RA and continuous RA were undertaken. The gene-based analysis of low RA identified two genes significantly associated with low RA: Forkhead Box J1 (*FOXJ1*) on chromosome 17, and Zinc Finger FYVE-type Containing 21 (*ZFYVE21*) on chromosome 14 (Supplemental Figure 3). The gene set analysis of continuous RA identified three genes: Copine 4 (*CPNE4*) and Chromosome 3 open reading frame 62 (*C3orf62*) on chromosome 3, and Renalase (*RNLS*) on chromosome 10 (Supplemental Figure 4).

### Genetic correlation between low RA and psychiatric phenotypes

There was preliminary evidence of genetic correlation between low RA and insomnia (r_g_=0·90, se=0·42, p=0·033), suggesting that the biology underlying low RA may be associated with the regulation of sleep (Table 1). However, on FDR correction this was not statistically significant. There were no significant genetic correlations identified between low RA and ADHD, anxiety, BD, MDD, mood instability, PTSD and schizophrenia.

**Table 1.** Genetic correlations between low relative amplitude and ADHD, anxiety, BD, MDD, mood instability, PTSD, schizophrenia and insomnia

| Phenotype | $r_g$ | se | z | p | p FDR corrected |
| --- | --- | --- | --- | --- | --- |
| ADHD | 0.3476 | 0.4541 | 0.7656 | 0.444 | 0.932 |
| Anxiety | -0.0037 | 0.4224 | -0.0088 | 0.993 | 1.000 |
| BD | -0.0626 | 0.1842 | -0.3396 | 0.7341 | 1.000 |
| MDD | 0.0045 | 0.2527 | 0.0178 | 0.9858 | 1.000 |
| Mood Instability | -0.1587 | 0.2713 | -0.5847 | 0.5587 | 0.939 |
| PTSD | 0.7396 | 0.6201 | 1.1926 | 0.233 | 0.713 |
| Schizophrenia | 0.1541 | 0.1353 | 1.139 | 0.2547 | 0.713 |
| Insomnia | 0.8996 | 0.4222 | 2.1308 | **0.0331** | 0.278 |

R<sub>g</sub> is the genetic correlation with low RA, se is the standard error of the correlation, z represents test statistic. p is the uncorrected p value and p FDR corrected has been adjusted for multiple testing.

### Association between PRS for low RA and affective disorder phenotypes

The findings of analyses assessing the association between low RA PRS and several mood disorder-related phenotypes are presented in Table 2. Positive associations were identified between increased PRS and mood instability at all PRS thresholds, with the exception of genome-wide significance (p<5×10^−8^). For MDD, small positive associations were found for the low RA PRS at the top three significance thresholds (OR=1·02-1·03), which remained significant after FDR correction (p=0·025-0·05). A positive association with neuroticism was found for the highest threshold (p=0·004, FDR adjusted p=0·021). However, other associations between the remaining PRS thresholds and neuroticism score were not significant.

**Table 2.** Associations between low RA PRS and psychiatric phenotypes

| PRS p threshold | Outcome<br>(Cases/Controls) | OR (95% CI) | p<br>uncorrected | p FDR<br>corrected |
| --- | --- | --- | --- | --- |
| $p < 5 \times 10^{-8}$ | BD<br>(406/37 699) | 0.99 (0.92, 1.06) | 0.748 | 0.785 |
| $p < 5 \times 10^{-5}$ | | 1.05 (0.98, 1.12) | 0.206 | 0.746 |
| $p < 0.01$ | | 1.03 (0.96, 1.11) | 0.355 | 0.746 |
| $p < 0.05$ | | 1.04 (0.96, 1.12) | 0.309 | 0.746 |
| $p < 0.1$ | | 1.02 (0.94, 1.10) | 0.617 | 0.785 |
| $p < 0.5$ | | 1.02 (0.93, 1.11) | 0.754 | 0.785 |
| $p < 5 \times 10^{-8}$ | MDD<br>(9 543/24 317) | 1.00 (0.99, 1.02) | 0.812 | 0.805 |
| $p < 5 \times 10^{-5}$ | | 1.00 (0.98, 1.02) | 0.966 | 0.805 |
| $p < 0.01$ | | 1.01 (0.99, 1.03) | 0.395 | 0.494 |
| $p < 0.05$ | | 1.02 (1.00, 1.04) | **0.03** | **0.05** |
| $p < 0.1$ | | 1.03 (1.01, 1.05) | **0.005** | **0.025** |
| $p < 0.5$ | | 1.03 (1.00, 1.05) | **0.021** | **0.05** |
| $p < 5 \times 10^{-8}$ | GAD<br>(2 587/23 564) | 0.97 (0.95, 1.00) | 0.092 | 0.3 |
| $p < 5 \times 10^{-5}$ | | 0.98 (0.96, 1.01) | 0.274 | 0.548 |
| $p < 0.01$ | | 0.99 (0.97, 1.02) | 0.729 | 0.729 |
| $p < 0.05$ | | 1.01 (0.98, 1.04) | 0.475 | 0.713 |
| $p < 0.1$ | | 1.03 (0.99, 1.06) | 0.1 | 0.3 |
| $p < 0.5$ | | 1.01 (0.97, 1.04) | 0.699 | 0.729 |
| $p < 5 \times 10^{-8}$ | Mood Instability<br>(78 710/91 248) | 1.00 (0.99, 1.01) | 0.913 | 0.94 |
| $p < 5 \times 10^{-5}$ | | 1.01 (1.00, 1.02) | **0.019** | **0.0096** |

| p<0.01 |  | 1.01 (1.01, 1.02) | **9.5x10<sup>-5</sup>** | **2.2x10<sup>-4</sup>** |
| --- | --- | --- | --- | --- |
| p<0.05 |  | 1.02 (1.01, 1.02) | **3.6x10<sup>-6</sup>** | **9.6x10<sup>-5</sup>** |
| p<0.1 |  | 1.01 (1.01, 1.02) | **8.3x10<sup>-5</sup>** | **5.9x10<sup>-4</sup>** |
| p<0.5 |  | 1.02 (1.01, 1.03) | **1.2x10<sup>-6</sup>** | **1.5x10<sup>-5</sup>** |
| PRS p threshold | Outcome | Beta (95% CI) | p<br>uncorrected | p FDR<br>corrected |
| p<5x10<sup>-8</sup> | Neuroticism score | -0.004 (-0.02, 0.007) | 0.456 | 0.399 |
| p<5x10<sup>-5</sup> | (140 504) | 0.01 (-0.003, 0.02) | 0.124 | 0.134 |
| p<0.01 |  | 0.01 (-0.002, 0.02) | 0.098 | 0.134 |
| p<0.05 |  | 0.01 (-0.005, 0.03) | 0.059 | 0.134 |
| p<0.1 |  | 0.01 (-0.003, 0.02) | 0.128 | 0.134 |
| p<0.5 |  | 0.02 (0.007, 0.04) | **0.004** | **0.021** |

## Discussion

We present the first large-scale genetic study of an objective measure of circadian rest-activity rhythmicity in humans, as well as the first study to examine how common risk SNPs for circadian disruption might be associated with mood disorder phenotypes. The primary GWAS of low RA identified three genome-wide significant SNPs within two independent loci; two of the SNPs highlighted were found to be in relatively high LD (r^2^=0·66-1·00) in many populations^44^ and are potentially tagging a single underlying functional variant influencing low RA. The secondary GWAS of RA as a continuous measure also identified five genome-wide significant SNPs at a single locus, within the *MEIS1* gene on chromosome 2. Again, the SNPs identified are in medium to high LD with each other (r^2^=0·39-1·00) in many populations.^45^

### Genes of interest

Of considerable interest is that one of the genes highlighted by GWAS was *NFASC*, a gene encoding the neurofascin protein.^46^ Neurofascin is an L1 family immunoglobulin cell adhesion molecule that interacts with several proteins to anchor voltage-gated Na^+^ channels to the intracellular skeleton in neurons.^46^ It is involved in neurite outgrowth and organization of axon initial segments (AIS) during early development.^46^ These AIS complexes (comprising neurofascin, ankyrin G (encoded by *ANK3*), gliomedin and betaIV spectrin) are important for the generation of action potentials and for the maintenance of neuronal integrity.^47^ Notably, polymorphisms in *ANK3* are robustly associated with BD.^48^ The direct binding of neurofascin to ankyrin G at the AIS therefore represents a potentially important biological link between circadian rhythmicity and BD. The *NFASC* SNPs identified by the GWAS were intronic variants and the *NFASC* transcript undergoes extensive alternative splicing with not all variants being functionally categorised.^46^ The precise influence these variants have on the *NFASC* gene is currently unclear.

One of the genome-wide significant SNPs from the primary GWAS is located within *SLC25A17*. This gene encodes a peroxisomal solute carrier membrane protein that transports several cofactors from the cytosol to the peroxisomal matrix.^49^ Variants of this gene have previously been associated with autism spectrum disorder and schizophrenia.^50^ The *SLC25A17* gene is also involved in adrenomyeloneuropathy,^51^ an inherited condition in which long chain fatty acids accumulate in the central nervous system (CNS) disrupting several brain functions.^52^ The SNP identified in the GWAS of low RA (rs9611417) associated with lower expression of *RANGAP1*, a GTPase activator protein involved in nuclear transport.^53^ *RANGAP1* is 439kb downstream of rs9611417 and shows relatively high expression in the brain (Supplemental Figure 6).

The GWAS of RA as a continuous measure highlighted SNPs within the *MEIS1* gene. This gene encodes a homeobox transcription factor (TF) protein crucial for the normal development of several tissues, including the CNS.^54,55^ The *MEIS1* gene is also associated with myeloid leukaemia^54^ and restless leg syndrome 7 (a sleep-wake disorder).^54,55^

### Gene-based analyses

The gene-based analysis of low RA identified two genes: *FOXJ1* on chromosome 17, and *ZFYVE21* on chromosome 14. *FOXJ1* encodes a forkhead TF protein which has a role in cell differentiation in respiratory, reproductive, immune, and CNS tissues. *FOXJ1* is required for the formation of cilia.^56^ In mouse models, *FOXJ1* was reported to be important for neurogenesis within the forebrain and olfactory bulb.^56^ The *ZFYVE21* gene encodes the zinc-finger FYVE-type containing 21 protein, involved in cell migration and adhesion.^57^ The potential involvement of this gene in the brain and circadian function is currently unclear.

For the continuous measure of RA, three genes were identified in the gene-based analysis: *CPNE4* and *C3orf62* (chromosome 3); and *RNLS* on (chromosome 10). The *CPNE4* gene encodes a calcium-dependent phospholipid binding protein involved in membrane trafficking and may be involved in intracellular calcium-mediated processes.^58^ Deletion of this gene has been associated with earlier age-of-onset of Alzheimer’s disease.^59^ Currently, the C3orf62 gene is an uncharacterised protein coding gene that has not been functionally annotated.^60^

RNLS encodes a flavin adenine dinucleotide-dependent amine oxidase, known as Renalase, an enzyme hormone secreted from the kidney into the bloodstream.^61^ Renalase is involved in mediating cardiac function and blood pressure by influencing heart rate and has been associated with hypertension, chronic kidney failure and type 1 diabetes prediction.^61–64^ It is worth noting that disrupted circadian rhythmicity has been associated with both diabetes and hypertension in several studies,^4,65,66^ and that RNLS has been associated with both treatment outcome and episode recurrence in BD.^67^

### Polygenic risk for low RA and psychiatric phenotypes

We found little evidence of genetic correlation between low RA and psychiatric phenotypes, even though disruption of circadian rhythmicity is a core feature of mood disorders and polymorphisms in circadian clock genes have been associated with BD within case-control studies.^68,69^ Further, as noted above, within the UK Biobank cohort we recently found that lower RA was associated with several adverse mental health outcomes.^24^ In the current study, we found some evidence for association between greater polygenic risk for low RA and both MDD and neuroticism. Across several PRS thresholds, we identified a more robust association between increasing PRS for low RA and the phenotype of mood instability. Mood instability is a common symptom that cuts across several psychiatric disorders.^70^ The possibility of a direct link between genetic loading for circadian disruption and the experience of dysregulated or unstable mood is therefore of considerable interest and merits further investigation.

### Limitations

Several limitations are acknowledged. UK Biobank may not be fully representative of the general UK population,^71^ with a possible under-representation of individuals with psychiatric disorders. The scores created using the genome-wide SNPs did not show any associations with lifetime mood disorder diagnoses, mood instability or neuroticism outcomes tested. As only 3 genome-wide significant SNPs were included in these analyses, they may be underpowered. The PRSs (at other thresholds) showed relatively small effects overall on the traits investigated and appeared to explain only a small proportion of the variance within the traits. In terms of phenotyping, the mood instability phenotype was a self-reported subjective measure that may be influenced by response bias. Further, we were unable to identify a direction of causality between RA and the psychiatric phenotypes: approaches to causality such as Mendelian randomisation (MR) were not possible due to the small number of genome-wide significant SNPs identified by both GWAS analyses.^72^

## Conclusions

Overall, our findings contribute substantial new knowledge on the genetic architecture of circadian rhythmicity and how this overlaps with mood disorder phenotypes, particularly mood instability. Several of the genetic variants identified are located within or close to genes which may have a role in the pathophysiology of mood disorders. It is hoped that these findings will act as stimulus for future work assessing the complex biological relationships between circadian function and psychiatric disorders.

## Acknowledgements

We thank all participants in the UK Biobank study. UK Biobank was established by the Wellcome Trust, Medical Research Council, Department of Health, Scottish Government and Northwest Regional Development Agency. UK Biobank has also had funding from the Welsh Assembly Government and the British Heart Foundation. Data collection was funded by UK Biobank. RJS is supported by a UKRI Innovation-HDR-UK Fellowship (MR/S003061/1). JW is supported by the JMAS Sim Fellowship for depression research from the Royal College of Physicians of Edinburgh (173558). AF is supported by an MRC Doctoral Training Programme Studentship at the University of Glasgow (MR/K501335/1). KJAJ is supported by an MRC Doctoral Training Programme Studentship at the Universities of Glasgow and Edinburgh. DJS acknowledges the support of the Brain and Behaviour Research Foundation (Independent Investigator Award 1930) and a Lister Prize Fellowship (173096). JC acknowledges the support of The Sackler Trust and is part of the Wellcome Trust funded Neuroimmunology of Mood and Alzheimer’s consortium that includes collaboration with GSK, Lundbeck, Pfizer and Janssen & Janssen. AD acknowledges the support of the NIHR Biomedical Research Centre, Oxford; and the British Heart Foundation Centre of Research Excellence at Oxford (RE/13/1/30181). The funders had no role in the design or analysis of this study, decision to publish, or preparation of the manuscript.

## Contributors

All authors contributed substantively to this work. DJS, MESB, JW, LML and AF were involved in study conception and design. CAW, JW, LML and AF were involved in data organisation and statistical analyses. DJS, MESB DML and CAW were involved in application to UK Biobank and data coordination. AF, LML and DJS drafted the report. All authors were involved in reviewing and editing of the manuscript and approved it. DJS, JW, LML and AF had full access to all the data in the study and take responsibility for the integrity and accuracy of analyses.

## Supplemental

**Supplemental Figure 1.**
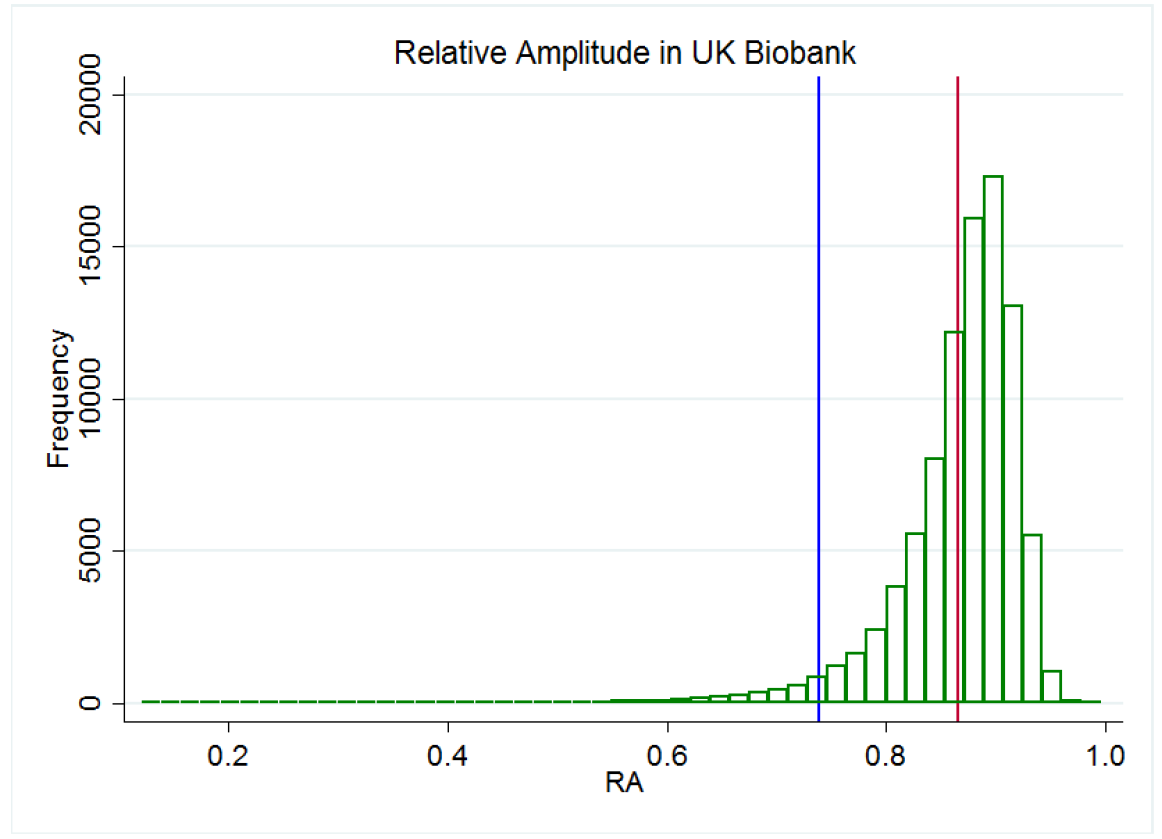
Relative amplitude histogram indicating (N=2 700 Cases and N=68 300 Controls) Red line represents mean value of RA. Blue line is two standard deviations from the mean, designating cases for use in the primary GWAS.

**Supplemental Figure 2.**
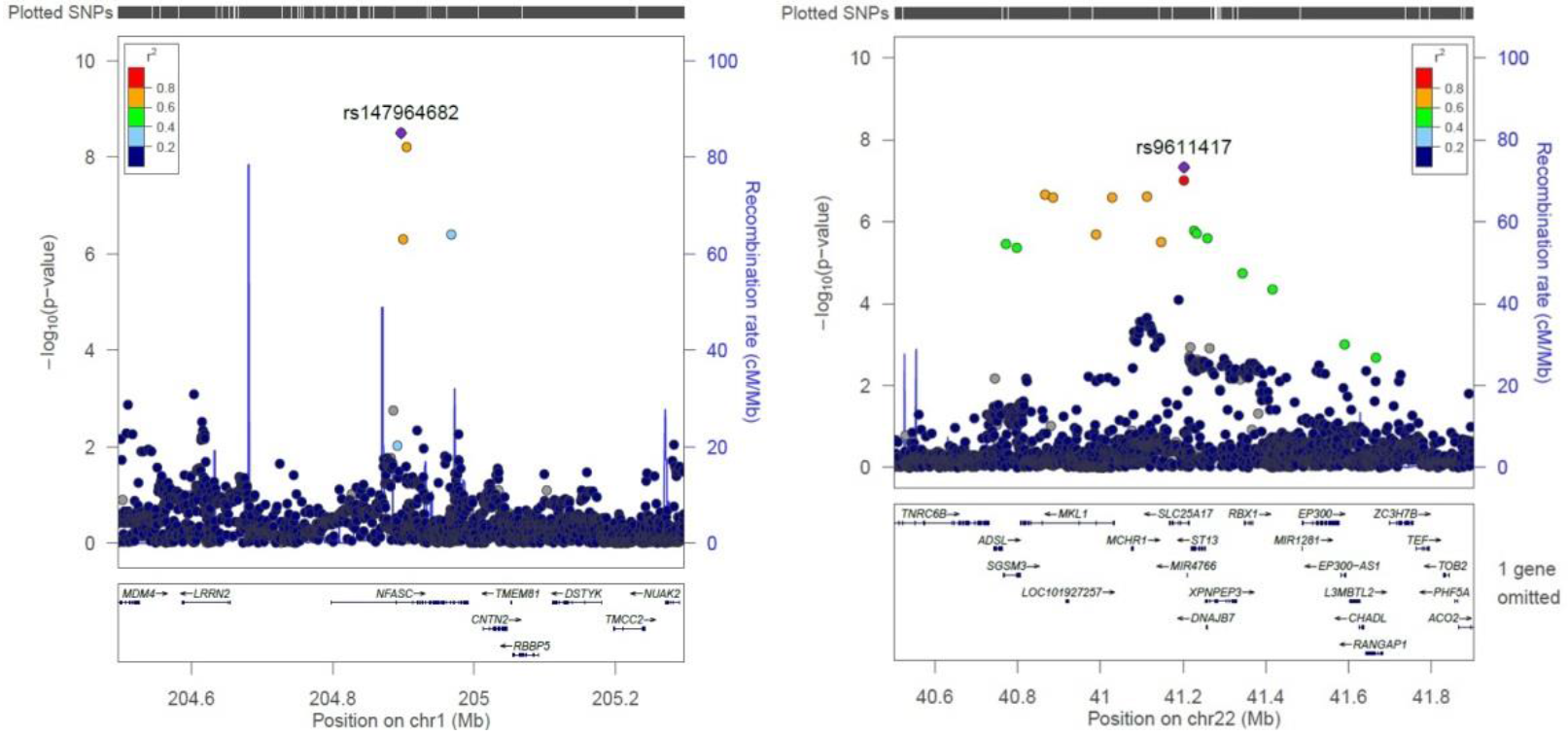
Regional plots of *NFASC* and *SLC25A17* Regional plots of SNPs produced by FUMA.^33^

**Supplemental Figure 3.**
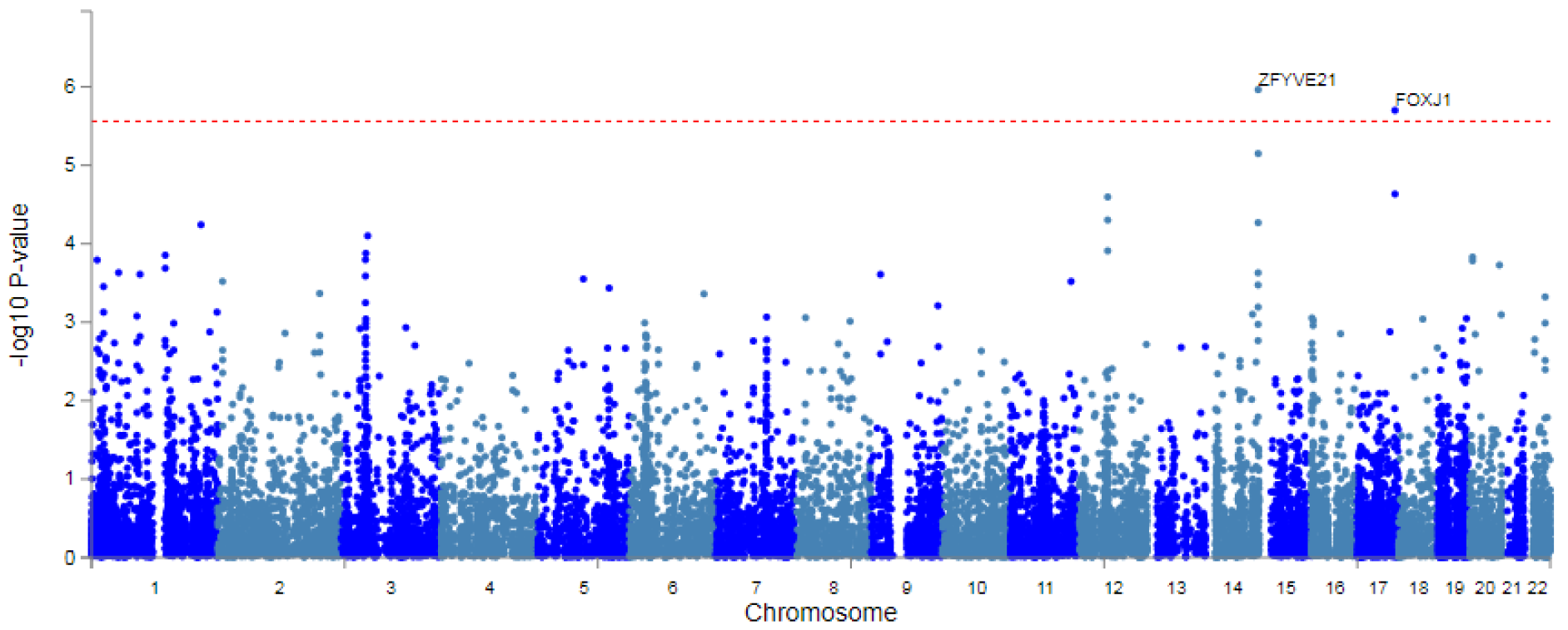
Low RA gene-based analysis Red line represents genome-wide significance (p<5×10^−8^).

**Supplemental Figure 4.**
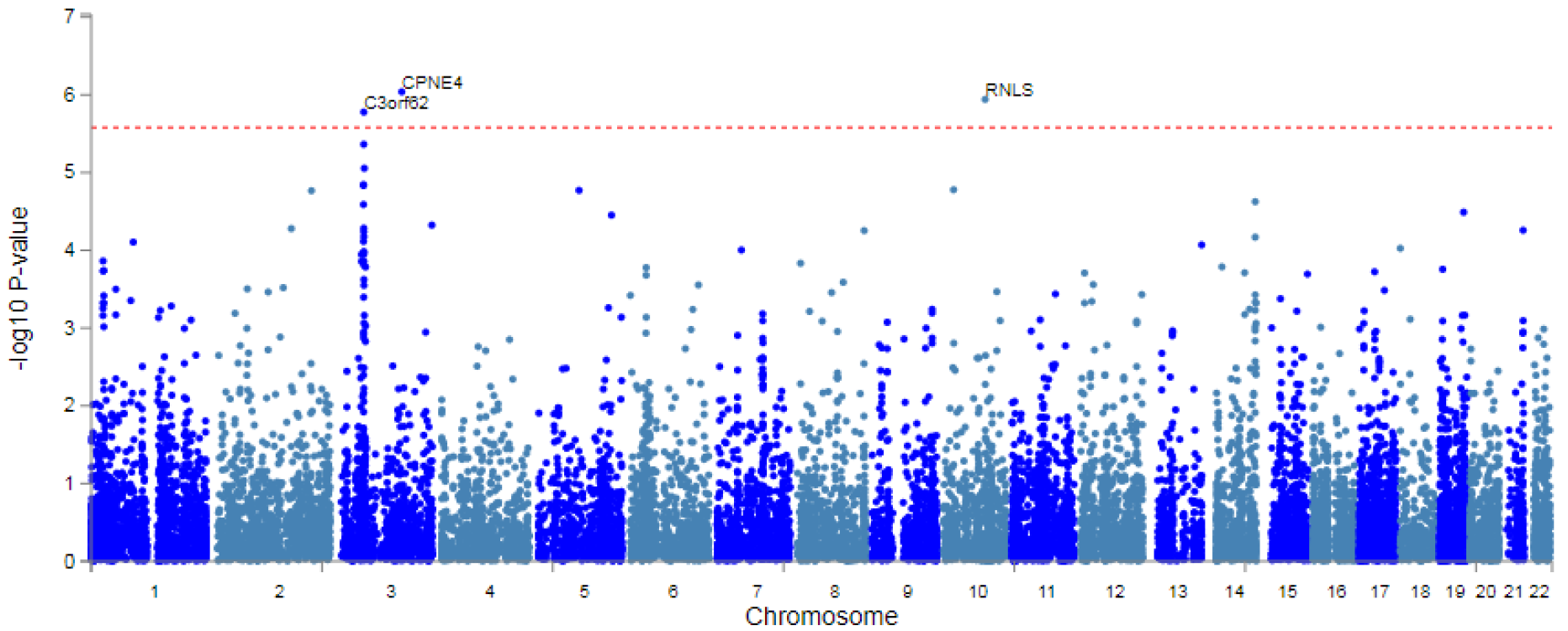
Continuous RA gene-based analysis Red line represents genome-wide significance (p<5×10^−8^).

**Supplemental Figure 5.**
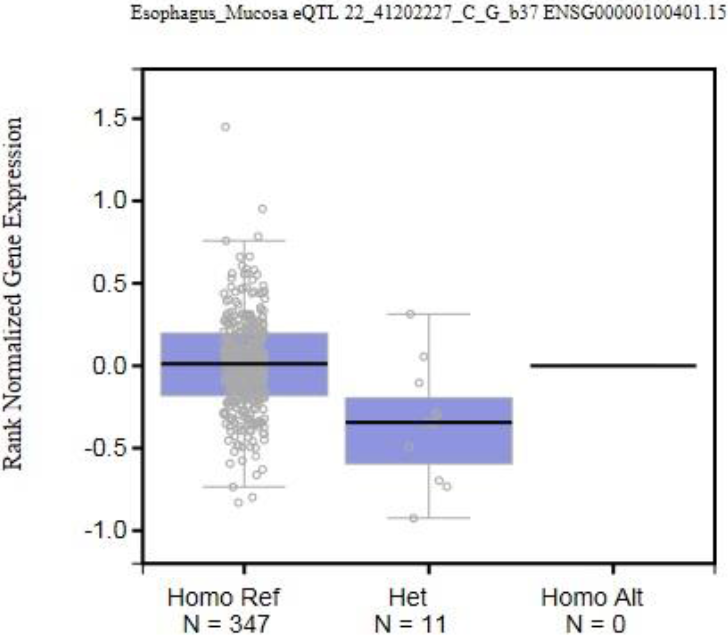
eQTLs of rs9611417 box plot Homo Ref: rs9611417 CC, Het: rs9611417 CG, Homo Alt: rs9611417 GG. Obtained from GTex portal.^32^

**Supplemental Figure 6.**
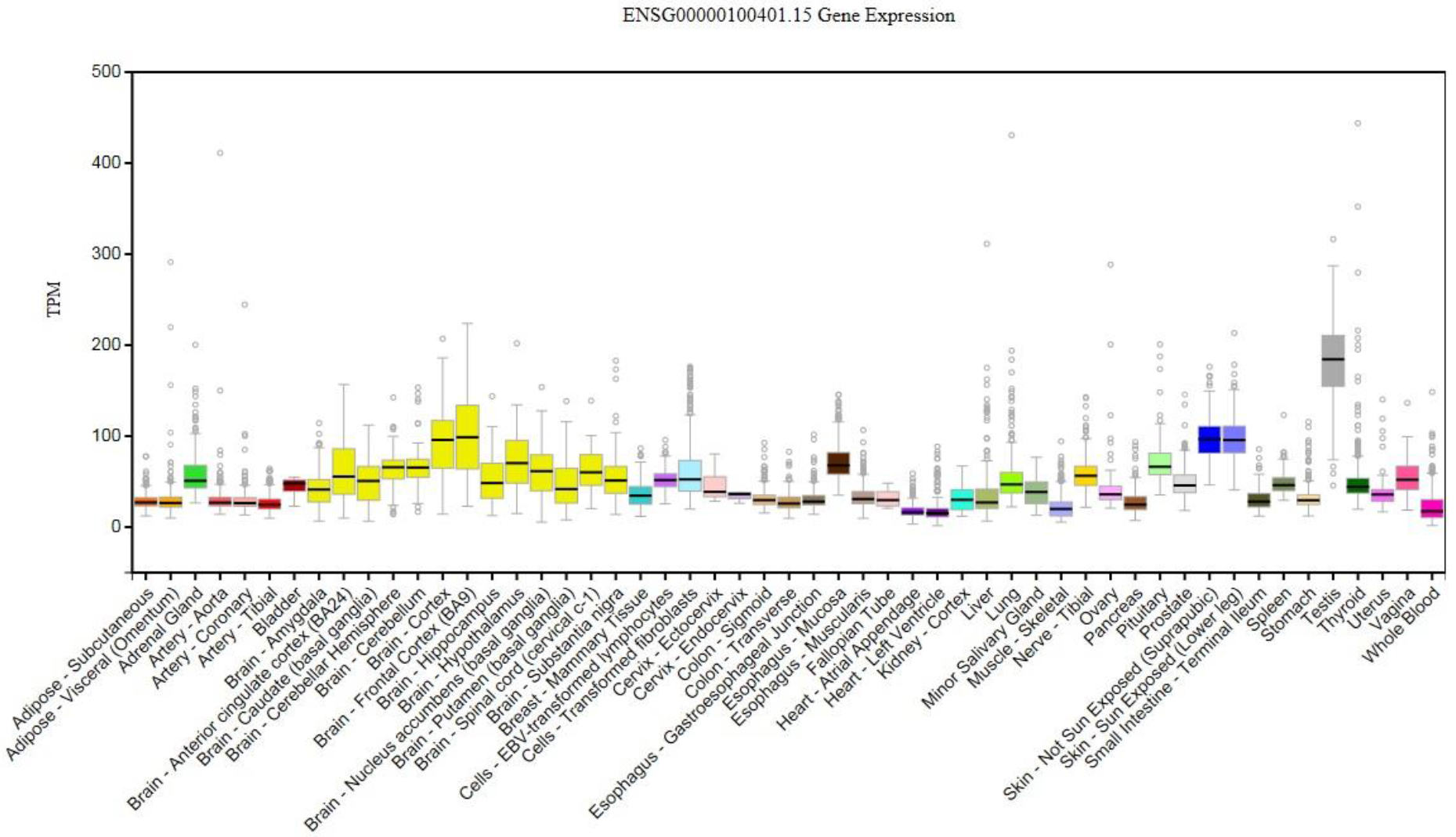
Tissue specific expression of *RANGAP1* Obtained from GTex portal.^32^

**Supplemental Table 1.** Genome-wide significant loci associated with low RA

| SNP | Chr | Position | A1/A2 | MAF | Beta | p | Nearby gene |
| --- | --- | --- | --- | --- | --- | --- | --- |
| rs147964682 | 1 | 204 896 804 | G/C | 0.012 | 0.584 | **3.179x10<sup>-9</sup>** | *NFASC* |
| rs146042826 | 1 | 204 904 528 | A/G | 0.013 | 0.568 | **6.17x10<sup>-9</sup>** | *NFASC* |
| rs9611417 | 22 | 41 202 227 | G/C | 0.012 | 0.5616 | **4.753x10<sup>-8</sup>** | *SLC25A17* |

**Supplemental Table 2.**
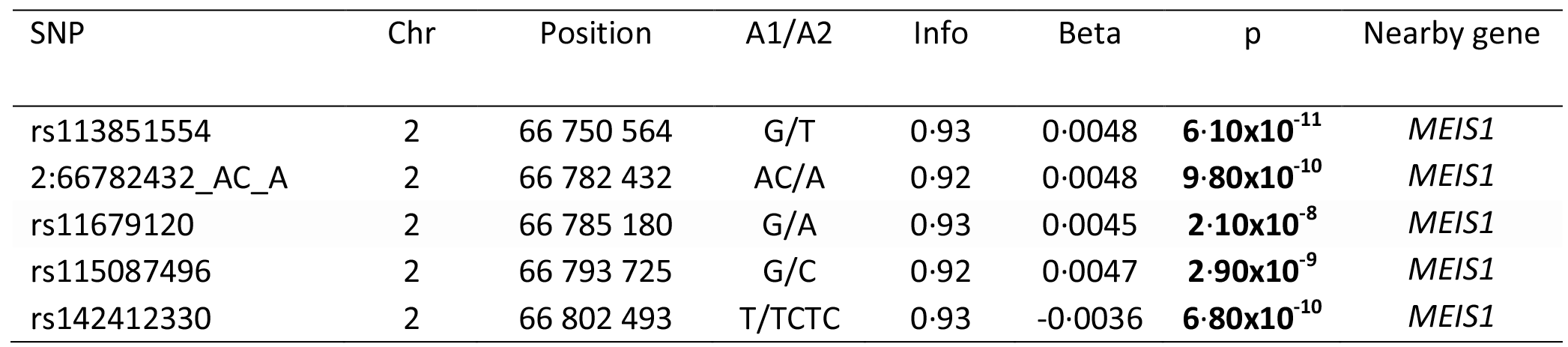
Genome-wide significant loci associated with continuous RA using BOLT-LMM

| SNP | Chr | Position | A1/A2 | Info | Beta | p | Nearby gene |
| --- | --- | --- | --- | --- | --- | --- | --- |
| rs113851554 | 2 | 66 750 564 | G/T | 0.93 | 0.0048 | **$6.10 \times 10^{-11}$** | *MEIS1* |
| 2:66782432_AC_A | 2 | 66 782 432 | AC/A | 0.92 | 0.0048 | **$9.80 \times 10^{-10}$** | *MEIS1* |
| rs11679120 | 2 | 66 785 180 | G/A | 0.93 | 0.0045 | **$2.10 \times 10^{-8}$** | *MEIS1* |
| rs115087496 | 2 | 66 793 725 | G/C | 0.92 | 0.0047 | **$2.90 \times 10^{-9}$** | *MEIS1* |
| rs142412330 | 2 | 66 802 493 | T/TCTC | 0.93 | -0.0036 | **$6.80 \times 10^{-10}$** | *MEIS1* |

